# Characterization of deep-sea viruses reveals their unexpected diversity and role in facilitating host metabolism of complex organic matter

**DOI:** 10.1101/2025.03.21.644689

**Authors:** Chong Wang, Rikuan Zheng, Chaomin Sun

## Abstract

Viruses exert a pervasive influence on biogeochemical cycles in deep-sea ecosystems. By integrating multi-omics and viral isolation data from four distinct deep-sea sediment sites, we uncover the diversity of DNA and RNA viral communities, which vary significantly across geographically separated cold seep and seamount environments. Host range predictions reveal both prokaryotic and eukaryotic communities are susceptible to viral infection. Metatranscriptomic analyses suggest viral functional genes are actively expressed, potentially enhancing host metabolism of complex organic matter (COM). Indeed, purified viruses from deep-sea sediments were shown to facilitate COM metabolism in certain deep-sea microorganisms, enabling the isolation of previously uncultured bacterial taxa. Notably, in these newly isolated bacteria, polysaccharides induced chronic virus release without cell lysis, a process that may assist host metabolism of polysaccharides. Our findings illuminate the diversity and roles of viral communities in distinct deep-sea sediments, underscoring their significant contribution to COM degradation and the deep-sea carbon cycle.

## Introduction

The seafloor is a diverse environment that ranges from the desert-like plains of deep-sea mud to the rich oases, which are present at cold seeps, hydrothermal vents, seamounts, trenches, and so on^1,2^. Cold seeps, typically located along continental margins, are regions where fluids rich in reduced compounds, such as methane, sulfide, and hydrocarbons, escape from the seafloor^3^. Seamounts are underwater mountains rising from the seafloor, often formed by volcanic activity^4^. These distinct geological features, though differing in their formation and physical characteristics, share a critical commonality: they are significant sites of enhanced organic matter supply and microbial-mediated biogeochemical cycling^5^. Understanding the intricate interplay between complex organic matter (COM), microbial communities, and viral dynamics in these deep-sea ecosystems is crucial for comprehending the global carbon cycle and the broader functioning of the marine biosphere.

Marine sediments represent the largest reservoir of organic carbon on Earth, accumulating material from diverse sources including primary production in the overlying water column, terrestrial runoff, cellular debris from lysis and mortality, and microbial exudates^6,7^. While surface sediment microbial communities are predominantly heterotrophic, relying on organic matter for carbon and energy^8–10^, continuous sedimentation over geological timescales results in the progressive burial and recalcitrance of this material^5^. Sedimentary organic matter comprises a complex mixture of biological macromolecules—carbohydrates, lipids, proteins, and nucleic acids—along with humic and fulvic acids^6,10,11^. Despite the extreme energy limitation and low metabolic rates characteristic of the deep subseafloor^1,5^, it is hypothesized that specialized microbial communities persist by utilizing these recalcitrant compounds^12^. Indeed, certain enriched deep-sea microbes exhibit the capacity to degrade biopolymers such as alginate, cellulose, pectin, and nucleic acids^13–15^. However, the precise mechanisms governing the microbial breakdown and utilization of COM in the deep sea remain poorly understood. In particular, the potential role of viruses in modulating these processes and shaping the fate of complex organic carbon remains largely unexplored.

Viruses are the most abundant and genetically diverse biological entities in the biosphere, exerting a profound influence on microbial communities and biogeochemical cycling^16^. In marine environments, viral abundance is estimated at 10 ^30, representing the second-largest biomass component after prokaryotes, despite their diminutive size^17,18^. Their ecological significance extends beyond host lysis, encompassing horizontal gene transfer and the modulation of host metabolism through auxiliary metabolic genes (AMGs)^19^. These AMGs can encode key metabolic enzymes, effectively rewiring cellular processes during infection and potentially influencing biogeochemical cycles. For instance, in peatland soils undergoing permafrost thaw in Sweden, virus-encoded glycoside hydrolases have been shown to play a significant role in the degradation of complex carbon polymers^20^, highlighting the capacity of viruses to directly impact carbon cycling in terrestrial ecosystems. While our understanding of viral diversity and activity in surface waters has grown considerably, knowledge of the deep sea remains limited, hampered by the logistical challenges of sample acquisition and analysis. Recent metagenomic surveys of cold seeps and seamounts, however, have begun to reveal the potential for viruses to shape deep-sea biogeochemistry. These studies have demonstrated that deep-sea viruses frequently carry AMGs associated with carbon, nitrogen, and sulfur metabolism, suggesting a capacity to augment prokaryotic host metabolism and influence elemental cycling in these unique and often energy-limited environments^21,22^. Of note, deep-sea ecosystems also harbor chronic viruses that exhibit a non-lytic release strategy: progeny phage particles are released from host cells via extrusion or budding without causing lysis^23–25^. These chronic viruses preserve host cell viability and potentially assist host metabolism^26^. Nevertheless, a comprehensive understanding of the diversity, lifestyles, and interactions with host cells of deep-sea DNA and RNA viral communities remains elusive.

In this study, we present a multi-omics investigation of four deep-sea sediments derived from the cold seep and seamount, integrating metagenomic, metatranscriptomic, DNA viromic, and RNA viromic datasets. We reveal distinct prokaryotic and viral community compositions across these contrasting environments, particularly in the newly formed cold seep. These findings highlight the central role of viruses in structuring deep-sea microbial communities, modulating their functional capacity, and influencing microbial-mediated degradation of COM. Furthermore, enrichment experiments with deep-sea viral fractions demonstrate that viral-encoded functions can augment host utilization of COM, providing a novel avenue for isolating and cultivating deep-sea difficult-to-culture microorganisms.

## Results and discussion

### Analysis of total organic carbon and prokaryotic community in different deep-sea sediments

To explore the diversity, host interaction, and ecological roles of viruses inhabiting deep-sea sediments, particularly their impact on host metabolism of complex organic matter (COM), we sequenced metagenomes, DNA and RNA viromes, and metatranscriptomes from four deep-sea sediment regions in the South China Sea. To account for the effect of deep-sea geographic features on microbial and viral communities, we collected sediment samples from geographically distinct locations: a declining cold seep vent (TPK), a site distant from the TPK (TPKF), a nascent cold seep vent (HMZ), and a seamount base (HSX) (Fig. 1a and Supplementary Data 1). For these four sediment samples, higher total organic carbon (TOC) content was found in sediments from the two cold seep vents, compared to sediments from the site distant from the seep and the seamount base (Fig. 1b). This disparity indicates a higher rate of microbial metabolism in cold seep vents, driven by the oxidation of reduced chemical compounds (e.g., methane and sulfide)^3^ and the enhanced availability of COM^13^. Our observation of elevated TOC levels further underscores the critical role of these chemosynthetically-driven ecosystems in deep-sea carbon cycling.

**Fig. 1.**
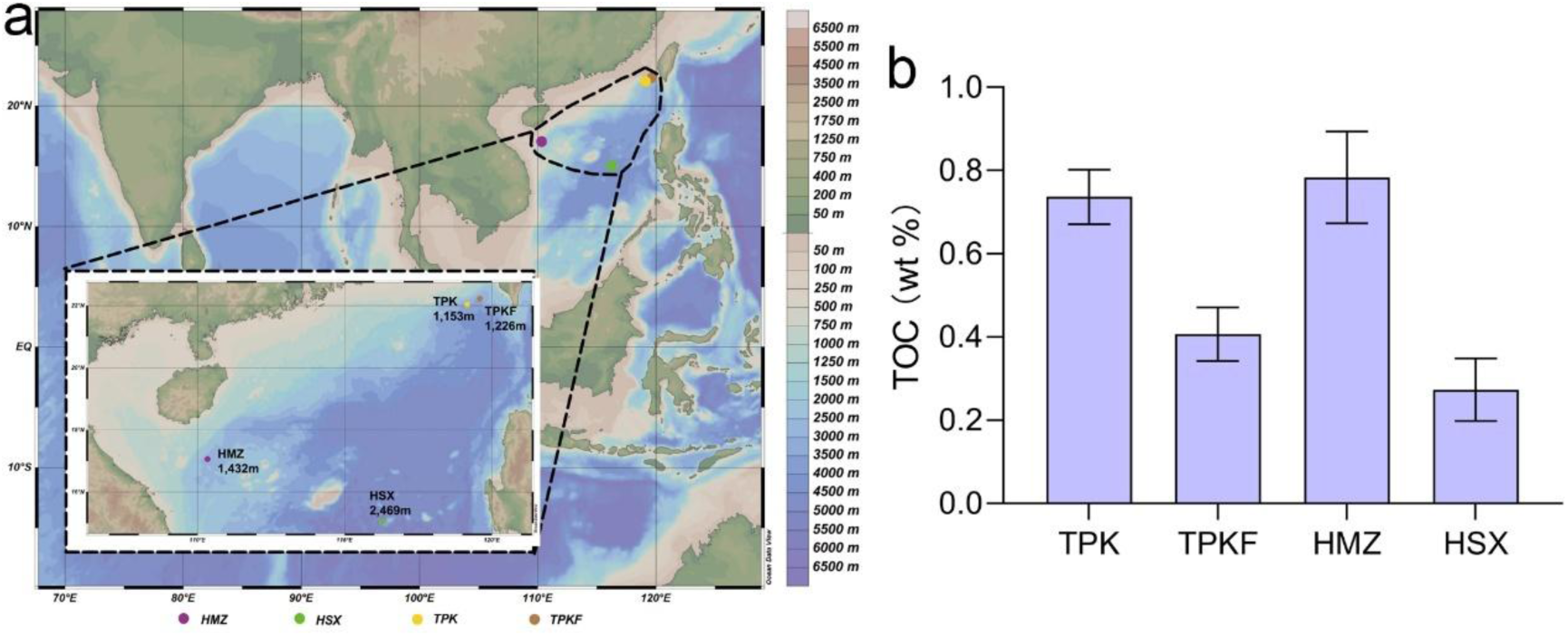
Geographic context and total organic carbon (TOC) in four deep-sea sediment samples. **a**, Geographic distribution of deep-sea sampling sites. Sampling sites and depths are colored according to sampling environments. This image was generated using Ocean Data View (ODV; Schlitzer, Reiner, *Ocean Data View*, https://odv.awi.de, 2021). **b**, Comparison of the TOC content of four deep-sea sampling sites mentioned in panel a. TPK, a declining cold seep vent; TPKF, a site distant from the TPK; HMZ, a nascent cold seep vent; HSX, a seamount base.

To assess the prokaryotic compositions in these four deep-sea sediments (TPK, TPKF, HMZ, and HSX), we performed metagenomic sequencing and analysis. *De novo* assembly and binning of metagenomes yielded 65 high- to medium-quality microbial metagenome-assembled genomes (MAGs; completeness ≥50%, contamination ≤10%)^27^. Clustering these MAGs at 95% average nucleotide identity (ANI) generated 58 bacterial and seven archaeal MAGs, representing species-level groups across 24 phyla (Supplementary Fig. 1 and Supplementary Data 2). The majority of bacterial MAGs belonged to dominant lineages, including *Pseudomonadota* (n = 10), *Bacteroidota* (n = 6), *Thermodesulfobacteriota* (n = 6), *Chloroflexota* (n = 5), *Planctomycetota* (n = 3), *Caldatribacteriota* (n = 3), *Omnitrophota* (n = 2), *Acidobacteriota* (n = 2), *Actinobacteriota* (n = 2), *Nitrospinota* (n = 1), *Myxococcota* (n = 1), and *Spirochaetota* (n = 1). Consistent with our findings, *Pseudomonadota* also dominate prokaryotic communities in diverse deep-sea sediments, including cold seeps, hydrothermal vents, and seamounts^22,28,29^. In addition to these common taxa, several MAGs affiliated with rare or uncultured lineages, such as *Patescibacteria* (n = 3), *Bipolaricaulota* (n = 1), *Cloacimonadota* (n = 1), *Fermentibacterota* (n = 1), *Krumholzibacteriota* (n = 1), *Latescibacterota* (n = 1), and *Methylomirabilota* (n = 1), were identified. Several MAGs could not be assigned to any known phylum, potentially representing novel lineages, and were predominantly recovered from HMZ, the newly formed cold seep sediment (Supplementary Data 2). Within the domain Archaea, MAGs were mainly affiliated with *Asgardarchaeota* (n = 2), *Thermoproteota* (n = 1), *Thermoplasmatota* (n = 1), and *Halobacteriota* (n = 3), with most originating from HMZ (Supplementary Data 2). These findings suggest that the deep sea, particularly newly formed cold seeps, harbors a diverse array of potentially novel microbial lineages.

**Supplementary Fig. 1.**
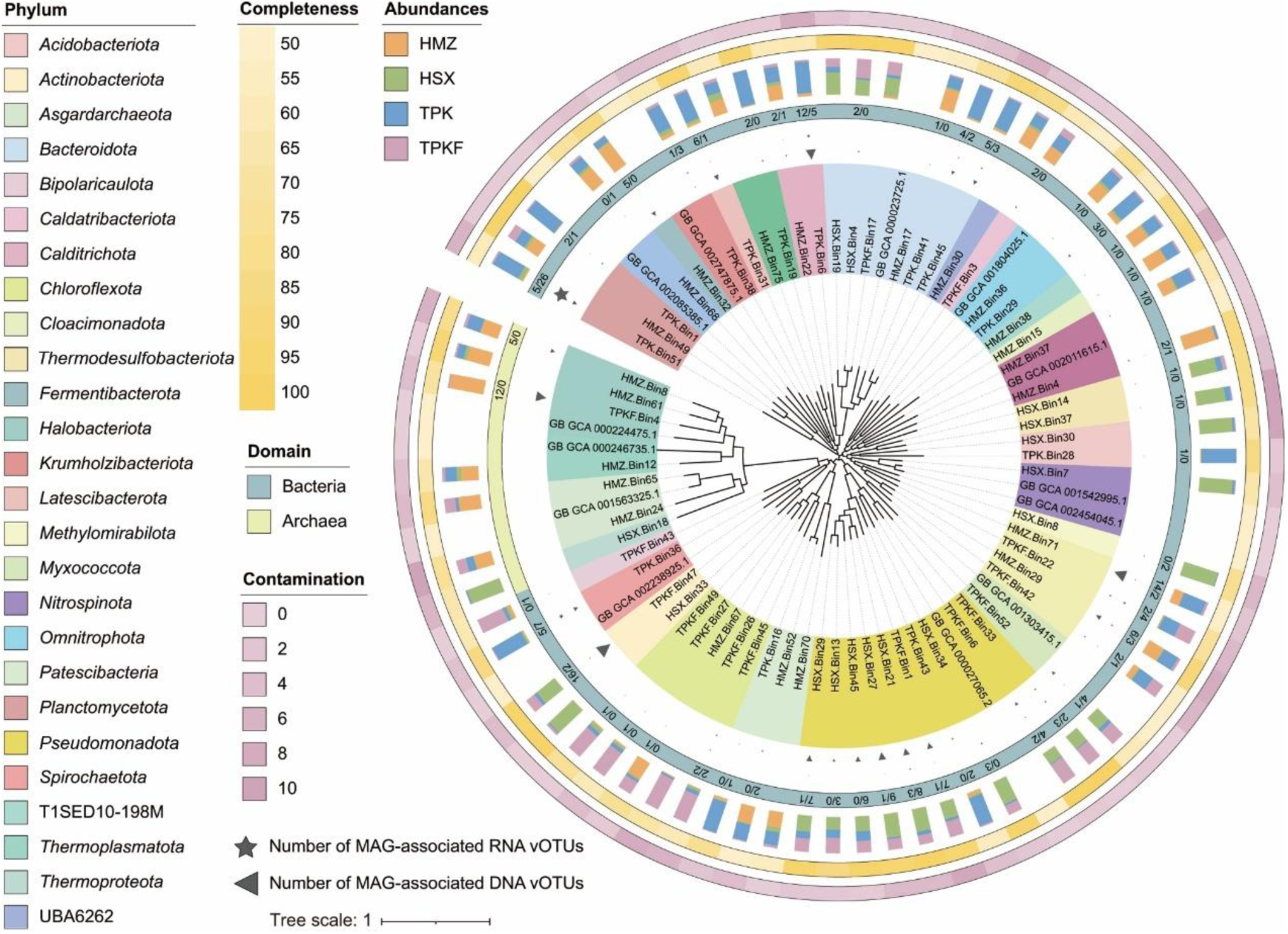
Maximum-likelihood phylogenetic tree of metagenome-assembled genomes (MAGs). Clades in the inner ring are colored according to their annotated phylum. In the middle ring, MAGs from Bacteria and Archaea are presented in gray-blue or green, respectively. Numbers within the central ring and the size of triangles and pentagons indicate the number of DNA and RNA viral operational taxonomic units (vOTUs) predicted to infect deep-sea MAGs, respectively. In the next outer ring, stacked columns indicate the relative abundance of MAGs across different sampling sites. The outermost two rings show the completeness and contamination levels of MAGs.

### Diverse viral morphologies revealed in deep-sea sediments

To investigate the presence and morphology of viruses in deep-sea sediments, we extracted viral particles from four above mentioned sediment samples (TPK, TPKF, HMZ, and HSX) and visualized them using transmission electron microscopy (TEM). This revealed a diverse assemblage of viral particles, demonstrating the widespread presence of viruses with varied morphologies in these deep-sea environments (Fig. 2). A few viruses have lipid envelopes or contain lipids as parts of internal vesicles (*Tectiviridae*) or the capsid (*Corticoviridae*) (Fig. 2a), and some viruses belonging to the *Globuloviridae* (Figs. 2b-c), which might exist in Archaea^30^. Two tailed virus-like morphotypes were observed at all deep-sea samples, corresponding to members of the *Myoviridae* (contractile tails) and *Siphoviridae* (long, flexible non-contractile tails) (Figs. 2d-e). A diverse array of filamentous (Fig. 2i) and rod-shaped (Figs. 2f, h, g) virus-like particles (VLPs) were also detected, which might belong to the *Inoviridae* and *Rudiviridae*. Notably, we observed a novel string-shaped virus in deep-sea sediments, characterized by a flexible, filamentous structure with repeating subunits (Fig. 2g). These structures, distinct from previously described viruses, suggest the potential discovery of a novel viral lineage with unique mechanisms of host infection and replication. Taken together, the ubiquitous presence of diverse viral assemblages, including numerous uncharacterized viral entities, across all four sediment samples underscores the deep sea as a crucial environment for investigating virus-host interactions and uncovering novel biological mechanisms adapted to these extreme conditions^31,32^. Further research into these unexplored viral communities holds the potential to reveal novel strategies for viral replication, host defense, and even inform the development of new antiviral therapies.

**Fig. 2.**
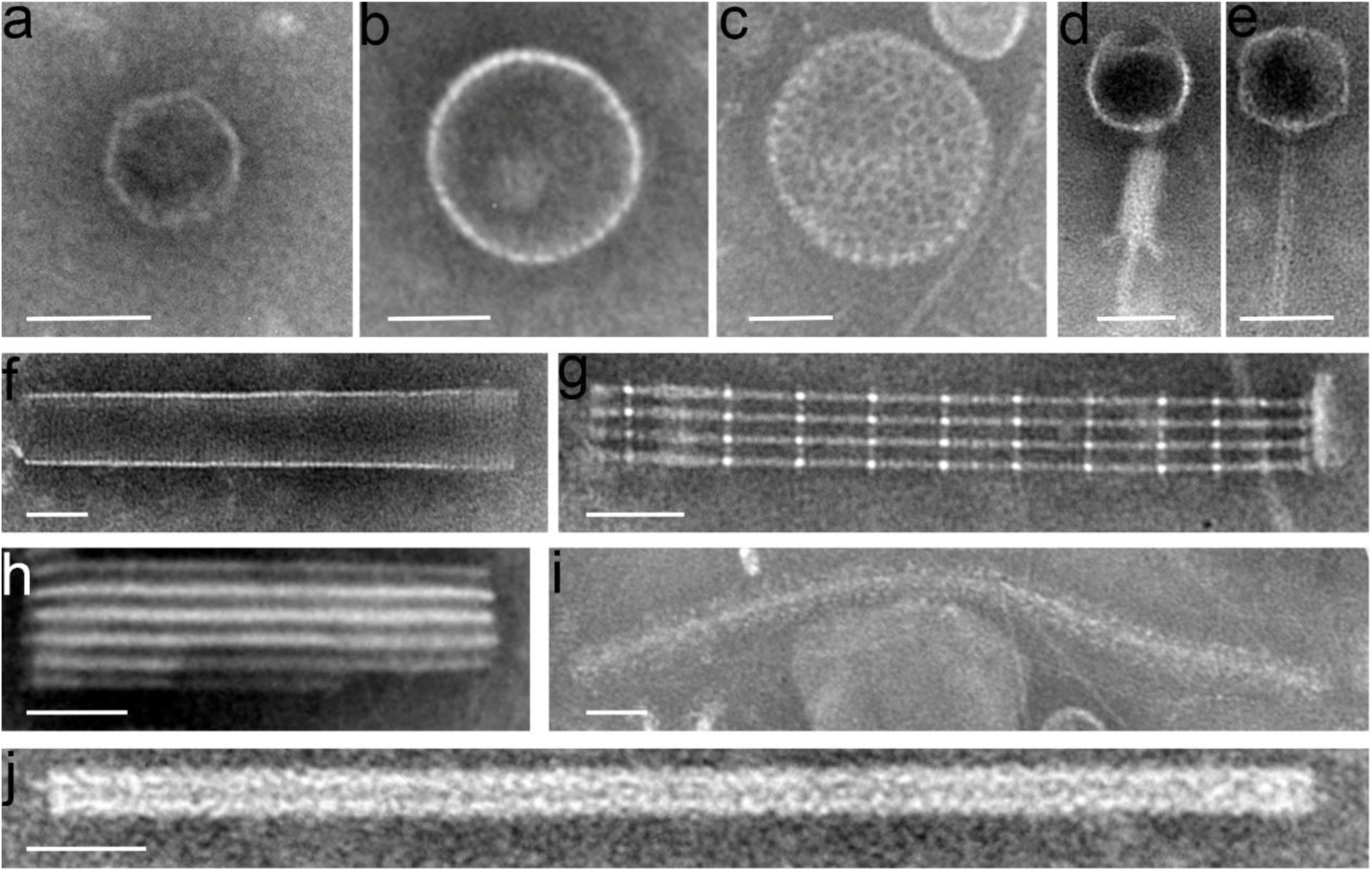
Transmission electron micrographs (TEM) showing the morphologies of virus-like particles in the deep-sea sediments. Representative images of virus-like particles are shown (panels a to j). Scale bars: 100 nm.

### Diversity, host interaction and lifestyle of DNA viral community in deep-sea sediments

To comprehensively characterize the diversity and abundance of DNA viruses in different deep-sea sediments, we performed DNA virome sequencing on four distinct sediment samples. Three pipelines were employed to identify viral sequences from the DNA virome datasets ^22^, resulting in 2,086 putative viral sequences (contigs ≥5 kb). To obtain larger viral assemblies, viral sequences were further binned using vRhyme v1.1.039. The resulting viral contigs and MAGs were then clustered at 95% identity and 85% coverage, generating 1,764 viral operational taxonomic units (vOTUs) (Supplementary Data 3). Taxonomic affiliations of the 1,764 vOTUs were determined by comparing predicted ORFs against the NCBI viral RefSeq database (v94) using the Last Common Ancestor algorithm. As shown in Fig. 3a and Supplementary Data 3, all 1,764 vOTUs could be taxonomically affiliated at the phylum level, with the majority assigned to *Uroviricota* (89.57%, n = 1,580; dsDNA tailed prokaryotic viruses), followed by *Nucleocytoviricota* (6.86%, n = 121; dsDNA NCLDVs), other phyla (1.47%, n = 26), and unclassified DNA viruses (2.10%, n = 37). To examine the viral community structures (at the family level) in four deep-sea sediments, clean reads from the DNA viromes were separately mapped to vOTUs to calculate reads per kilobase per million mapped reads (RPKM) values for each vOTU. In general, the abundance of single-stranded DNA (ssDNA) viruses was substantially lower than that of double-stranded DNA (dsDNA) viruses across all four deep-sea sediment samples (Fig. 3b and Supplementary Data 4). The newly formed cold seep (HMZ) exhibited lower DNA virus abundance compared to the gradually declining cold seep and seamount sediments, including *Poxviridae*, *Phycodnaviridae*, *Siphoviridae*, *Myoviridae*, and unclassified *Uroviricota*, suggesting that a mature viral-host interaction ecosystem has not yet been established at HMZ. We found that some dsDNA viruses, belonging to nucleocytoplasmic large DNA virus (NCLDV) and carrying NCLDV marker genes, were present in all four deep-sea sediment samples (Fig. 3b). However, most of their genomes were smaller than 70 kb, the known lower size limit for NCLDV genomes^33^, with only one reaching 81.8 kb, possibly due to incomplete assembly. The absence of known host contamination within these vOTUs suggests that they may represent authentic NCLDV genomes. It has been widely reported that NCLDVs readily exchange genetic material with hosts, prokaryotes, and bacteriophages^34^, leading to the acquisition of NCLDV marker genes by certain phages and their subsequent classification as members of the *Nucleocytoviricota*.

**Fig. 3.**
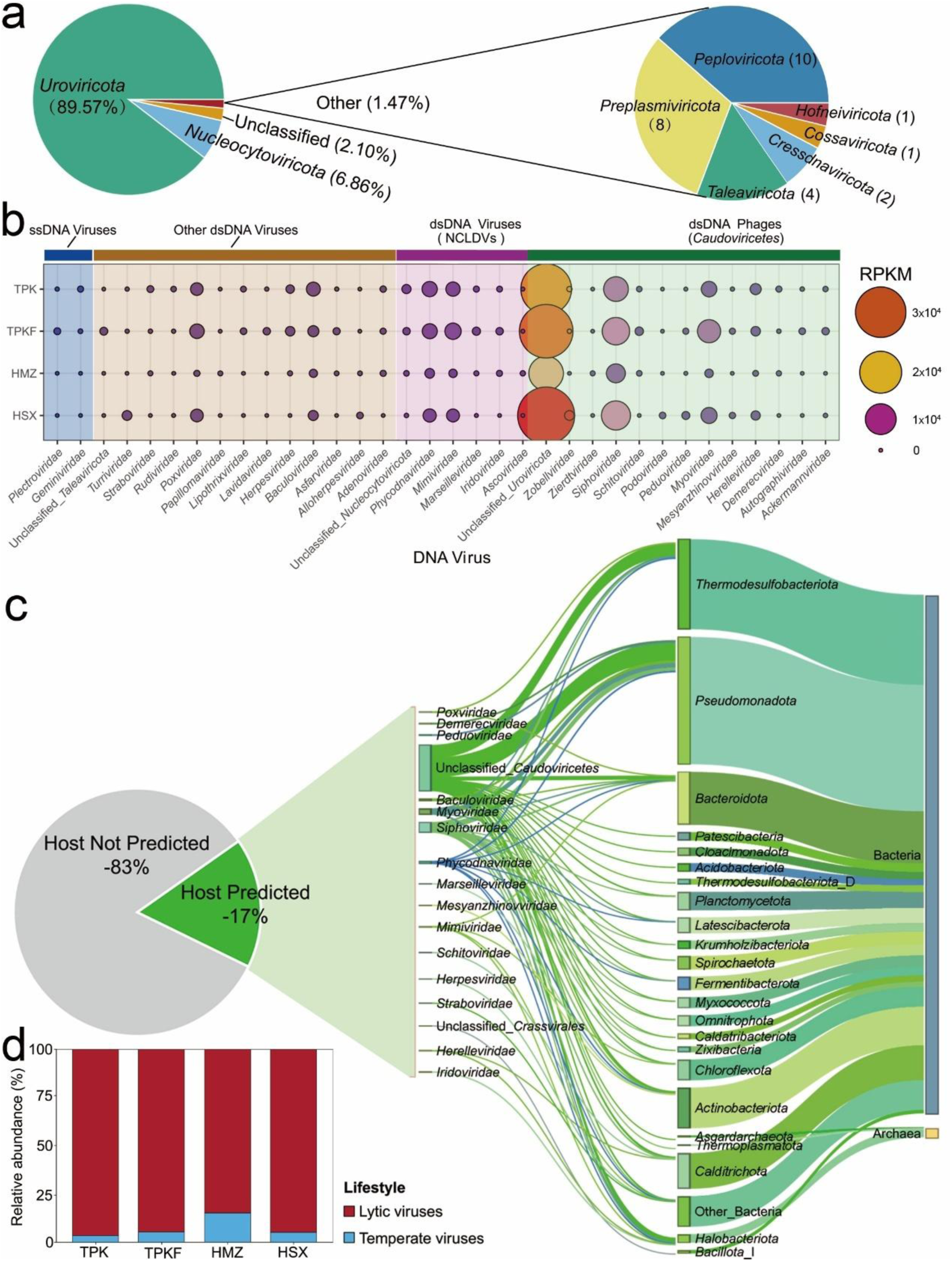
Community structure of deep-sea DNA viruses. **a**, Relative percentage of viral operational taxonomic units (vOTUs) at the phylum level. **b**, Relative abundance of DNA viruses at the family level in four deep-sea sediment samples (TPK, TPKF, HMZ, and HSX) derived from DNA virome data. **c**, Predicted virus-host linkages. The percentage and taxonomy of vOTUs for which a host was predicted are shown on the left; the taxonomy of predicted hosts is shown on the right. **d**, Predicted lifestyles of vOTUs derived from DNA virome data. Source data are provided as a Supplementary Data file.

To explore DNA virus-host interactions in deep-sea sediments, we predicted potential hosts for the 2,086 vOTUs using a combination of four bioinformatic approaches: sequence similarity, oligonucleotide frequencies, tRNA sequences, and CRISPR spacers^35^. To predict virus-host connections, we used a combination of host databases to infer virus-host linkages, including the MAGs binned from four deep-sea sediment samples in this study and cold seep samples from our previous study^36^. Consequently, 17% (n = 354) of these deep-sea vOTUs were assigned putative hosts, with 138, 69, 96, and 51 vOTUs predicted to infect hosts in the TPK, TPKF, HMZ, and HSX sediments, respectively (Fig. 3c and Supplementary Data 5). The substantial proportion of DNA viruses linked to four deep-sea sediment MAGs reveals a complex and pervasive network of virus-host interactions, suggesting a key role in shaping microbial community structure and biogeochemical cycling in the deep-sea benthos (Supplementary Fig. 1). Consistent with previous observations^35,37^, these vOTUs were predicted to have narrow host ranges, typically infecting a single host species. Predicted prokaryotic hosts spanned 3 archaeal and 19 bacterial phyla, with *Thermodesulfobacteriota* (33.9% of virus-host pairs), *Pseudomonadota* (11.0%), *Bacteroidota* (11.0%), and *Halobacteriota* (9.0%) being the most frequently predicted (Fig. 3c and Supplementary Data 6). Notably, viruses in the HMZ cold seep were predicted to infect archaeal hosts comprising 53.5% of the total predicted archaeal hosts, suggesting active virus-archaea interactions and associated metabolic activity in this nascent cold seep ecosystem. These findings suggest considerable divergence in viral communities across different deep-sea habitats. In addition, several families of NCLDV (*Phycodnaviridae*, *Mimiviridae*, *Marseilleviridae*, and *Iridoviridae*) were predicted to infect bacterial hosts (Fig. 3c). This highlights potential limitations in our current understanding and taxonomic classification of viruses carrying NCLDV marker genes, underscoring the need for isolation and cultivation to accurately determine their taxonomic placement and host range. Analogous to the recent, dramatic expansion of the phylum *Lenarviricota*—now encompassing both bacterial and related eukaryotic viruses—further investigation may reveal unexpected diversity and evolutionary relationships among NCLDVs^38^.

Moreover, the lifestyles of 1,764 vOTUs were predicted based on the presence of lysogeny-specific features, including genes encoding integrases, recombinases, and excisionases, and/or their genomic location within host chromosomes. Consequently, the vast majority (n = 1,465) of vOTUs were predicted to be lysogenic, whereas only a small minority (n = 42) were classified as temperate viruses (Supplementary Data 7). Based on abundances determined by read mapping, the relative abundance of temperate viruses in each sample was calculated for DNA virome datasets. As shown in Fig. 3d, the proportion of temperate phages was markedly higher in HMZ sediments (15.2%) compared to the other three deep-sea sediment samples (3.4%, 5.3%, and 5.1% for TPK, TPKF, and HSX, respectively). The prevalence of temperate viruses in these sediments, while potentially underestimated due to limitations in viral genome assembly and annotation that may obscure lysogeny-specific genes and flanking host sequences, aligns with observations from other deep-sea environments, including deep water from the South China Sea and diffuse-flow hydrothermal vents^39^. This suggests that temperate DNA viruses are particularly important in this nascent vent environment, where they may enhance host fitness, mediate horizontal gene transfer, modulate host lifecycles, and influence microbial community structure^40,41^, potentially contributing to host survival and stability in these challenging conditions^22^.

### Diversity, host interaction and lifestyle of RNA viral community in deep-sea sediments

While both DNA and RNA viruses play key roles in regulating biogeochemical cycles within marine ecosystems^42^, the deep-sea RNA virome, particularly in seamount and cold seep sediments, remains largely unexplored^43^. Therefore, we performed RNA virome sequencing on these four distinct sediment samples. Three pipelines were employed to identify RNA viral sequences from the RNA virome datasets, resulting in 1,021 putative RNA viral sequences (contigs ≥2 kb). The resulting viral contigs and MAGs were then clustered at 95% identity and 85% coverage, generating 767 viral operational taxonomic units (vOTUs) (Supplementary Data 8). Taxonomic analysis of 767 vOTUs, based on predicted ORFs compared against the NCBI viral RefSeq database (v94; Last Common Ancestor algorithm), revealed phylum-level affiliations for all vOTUs (Fig. 4a, Supplementary Data 8). The majority were assigned to *Kitrinoviricota* (47.33%, n = 363), followed by *Pisuviricota* (19.3%, n = 148), *Lenarviricota* (11.47%, n = 88), *Duplornaviricota* (10.43%, n = 80), *Negarnaviricota* (7.82%, n = 60), and *Artverviricota* (3.65%, n = 28). Community analysis, using the RPKM values, showed a substantial predominance of single-stranded RNA (ssRNA) viruses over double-stranded RNA (dsRNA) viruses across all sediments (Figs. 4b-c), in contrast to DNA viruses. The vOTUs that matched the known viral sequences were taxonomically assigned to 69 families of RNA viruses (Supplementary Data 9). This substantial proportion underscores the remarkable diversity of RNA viruses within deep-sea sediments, echoing global trends of high novelty^44^ and suggesting that these underexplored environments are reservoirs of novel viral lineages with potentially unique ecological functions. While RNA viral communities in all sediment samples were dominated by *Tymoviridae*, *Solspiviridae*, and *Steitzviridae*, the newly formed cold seep (HMZ) sediment exhibited a striking enrichment of *Xinmoviridae* (Fig. 4b). Given that *Xinmoviridae* are known to infect eukaryotic hosts^45^, this suggests a disproportionately strong virus-eukaryote interaction network within the HMZ region compared to virus-prokaryote interactions. This finding highlights the potential for eukaryotic microorganisms to play a significant, yet often overlooked, role in deep-sea viral ecology and biogeochemical cycling within deep-sea sediment environments.

**Fig. 4.**
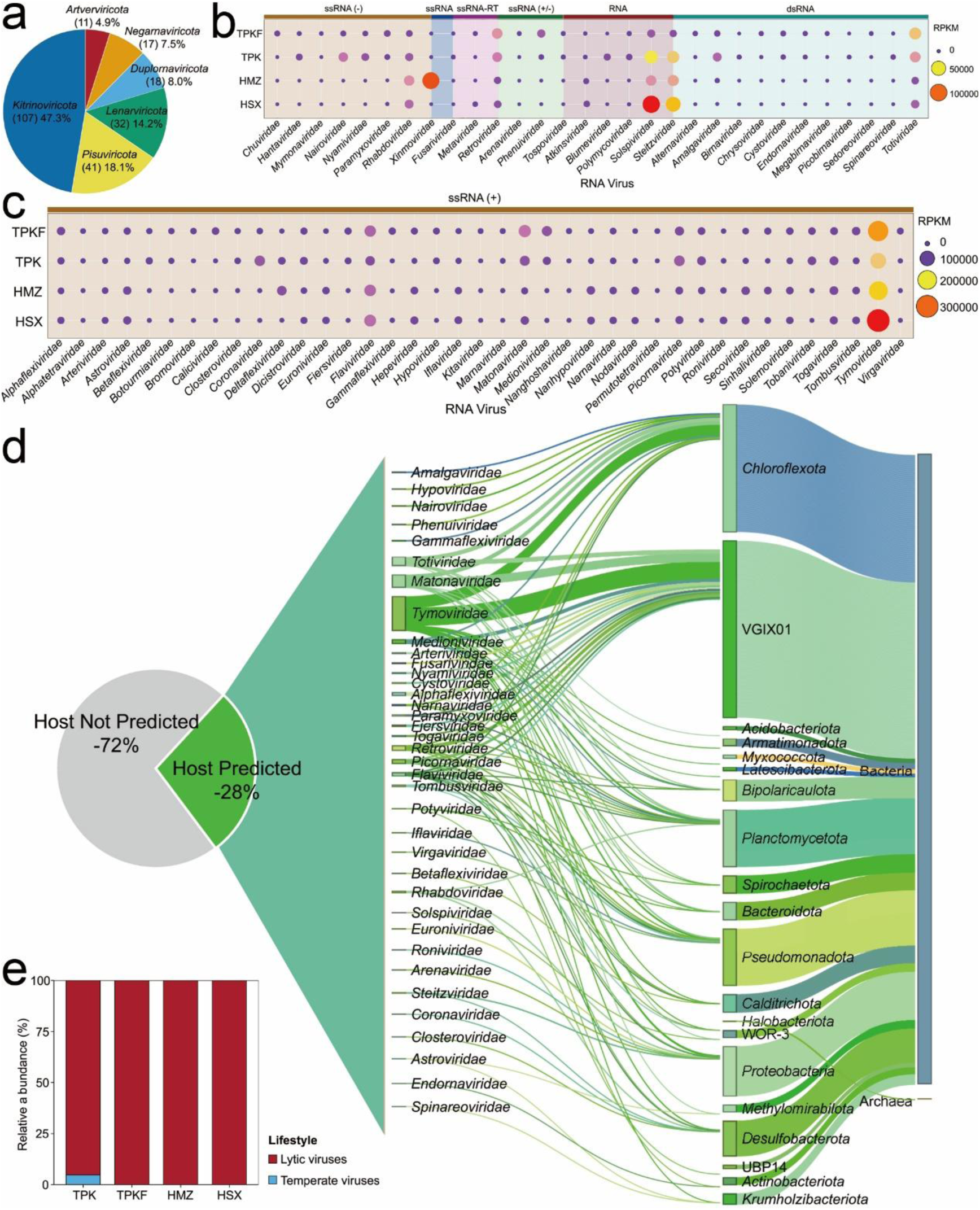
Community structure of deep-sea RNA viruses. **a**, Relative percentage of viral operational taxonomic units (vOTUs) at the phylum level. **b**, Relative abundance of RNA viruses at the family level in four deep-sea sediment samples (TPK, TPKF, HMZ, and HSX) derived from RNA virome data. **c**, Predicted virus-host linkages. The percentage and taxonomy of vOTUs for which a host was predicted are shown on the left; the taxonomy of predicted hosts is shown on the right. **d**, Predicted lifestyles for vOTUs derived from RNA virome data. Source data are provided as a Supplementary Data file.

To elucidate RNA virus-host interactions within these deep-sea sediments, predicted potential hosts for 1021 RNA vOTUs. This analysis assigned putative hosts to 28% (n = 285) of the vOTUs. Predicted prokaryotic hosts spanned 1 archaeal and 15 bacterial phyla, with *Chloroflexota* (18.2% of virus-host pairs), *Pseudomonadota* (13.7%), *Planctomycetota* (11.2%), and *Thermodesulfobacteriota* (7.0% of virus-host pairs) being the most frequently predicted (Fig. 4d and Supplementary Data 10-11). This distribution of predicted hosts largely mirrors that observed for viruses in deep-sea sediments, suggesting a conserved pattern of DNA and RNA viral predation across diverse prokaryotic lineages within these deep-sea environments. Notably, while 119, 151, and 14 vOTUs were predicted to infect prokaryotic hosts in TPK, TPKF, and HSX sediments, respectively, only a single RNA vOTU in the newly-formed HMZ sediment was predicted to infect a prokaryote (Supplementary Data 10). These data revealed that the eukaryotic viruses accounted a large proportion of the classified RNA viruses in the KMZ sediment. The overwhelming bias towards eukaryotic hosts in HMZ suggests a strong viral influence on eukaryotic microbial community structure and function within this unique, chemosynthetically-driven ecosystem. Moreover, the vast majority (n = 178) of vOTUs were predicted to be lysogenic, whereas only a single RNA vOTU was classified as temperate viruses (Supplementary Data 12). Based on abundances determined by read mapping, the relative abundance of temperate viruses in each sample was calculated for RNA virome datasets. Temperate phages comprised 4.7% of the RNA viral community in TPK sediments, but were notably absent from the other three sediment samples (Fig. 4e). Taken together, our findings suggest that despite their lower relative abundance compared to DNA viruses, RNA viruses in deep-sea sediments exhibit high diversity, a predominantly lytic lifestyle, and a strong propensity to infect eukaryotic hosts^32^. This combination of factors implies a potentially significant and previously underappreciated role for RNA viruses in structuring deep-sea microbial communities, particularly with respect to shaping eukaryotic population dynamics and influencing the flow of carbon and energy through deep-sea ecosystems.

### Potential impacts of viral functional genes on the degradation of COM

Viruses can reshape host metabolism through the expression of virus-encoded auxiliary metabolic genes (AMGs)^21,22^. Here, we broaden the concept of “viral functional genes” by defining them as both AMGs and previously unannotated viral genes with potentially significant impacts on host metabolism. To elucidate the potential impacts of viral functional genes on host metabolism during anaerobic macromolecule degradation, we functionally annotated genes from the above DNA and RNA vOTUs using the NR, Pfam, KEGG, and CAZy databases. Overall, metagenomic analysis of prokaryotic DNA viruses across four deep-sea sediments revealed a diverse repertoire of functional genes related to the metabolism of COM, including polysaccharides, proteins, and nucleic acids, with glycoside hydrolases (GH), and glycosyltransferase (GT), carbohydrate-binding module (CBM), exonuclease, HNH endonuclease, and peptidases being particularly prevalent (Fig. 5a). In contrast, prokaryotic RNA viruses harbored markedly fewer such genes, with none detected in the recently formed cold seep sediment HMZ (Fig. 5b), may be a consequence of their lower abundance and smaller genome size compared to DNA viruses. However, metatranscriptomic data revealed distinct expression patterns, with DNA viral functional genes more highly expressed in TPK and TPKF sediments, while RNA viral genes dominated expression in HMZ and HSX sediments (Figs. 5c-d and Supplementary Data 13). These findings suggest that both DNA and RNA viruses in the deep sea possess the capacity to augment prokaryotic host metabolism of COM, consistent with previous reports of abundant polysaccharide-degrading AMGs in deep-sea viruses^21,46^. Given that microorganisms associated with cold seep sediments are known to mediate carbohydrate degradation reactions essential for processing detrital organic matter from the water column^8,47^, these viral functional genes may play a critical role in facilitating host metabolism of COM, thereby influencing biogeochemical cycling in these unique ecosystems.

**Fig. 5.**
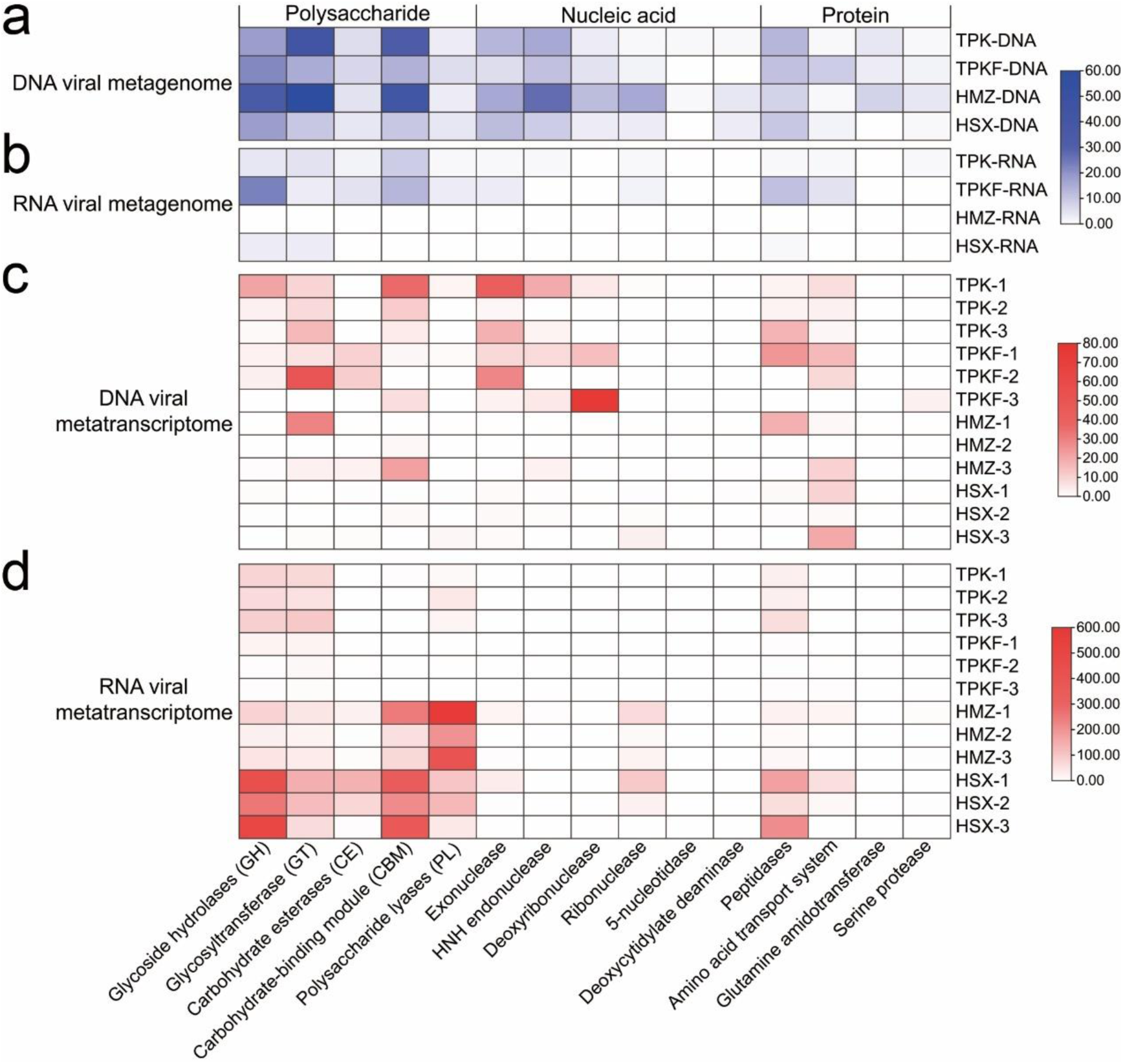
Viral functional genes and transcripts related to complex organic matter metabolism in deep-sea sediments. **a, b**, Relative abundance of functional genes within DNA and RNA viral metagenomes. **c, d**, Metatranscriptomic analysis quantified transcript abundance of viral functional genes (DNA and RNA) as transcripts per million (TPM).

### Purified viruses facilitate the enrichment and cultivation of deep-sea difficult-to-culture microorganisms

Given our findings that deep-sea DNA and RNA viruses harbor functional genes related to the degradation of COM, suggesting a potential role in augmenting host metabolism, we hypothesized that these viruses could influence microbial communities in deep-sea sediments. To test this, we extracted viral fractions from four distinct deep-sea sediment samples and used them to inoculate culture media containing COM and sediment samples, aiming to enrich novel microorganisms. To understand which bacterial taxa are present within the samples in the presence or absence of deep-sea viruses, we performed amplicon sequencing. The amplicon results showed that deep-sea viruses could effectively enrich difficult-to-culture microorganisms (including *Cyanobacteriota*, *Chloroflexota*, *Planctomycetota*, *Caldatribacteriota*, and *Acidobacteriota* bacteria) in four deep-sea sediments (TPK, TPKF, HMZ, and HSX) (Fig. 6a). Notably, the HMZ sediment sample exhibited the highest relative abundance of these enriched, previously unculturable microorganisms. We continuously cultured the enriched samples in the inorganic medium supplemented with polysaccharide and deep-sea viruses, and eventually obtained seven novel pure isolates (including *Lentisphaerota* strain WC36 (Supplementary Fig. 2a); *Planctomycetota* strain ZRK34 (Supplementary Fig. 2b); *Verrucomicrobiota* strain ZRK36 (Supplementary Fig. 2c); *Fusobacteriota* strain ZRK30 (Supplementary Fig. 2d); *Mycoplasmatota* strain zrk7; two *Bacillota* strains zrk8 and zrk18) and two enrichments (one is predominant by *Chloroflexota* strain ZRK35, and the other is predominant by *Cyanobacteriota* strain ZRK38) (Fig. 6b).

**Fig. 6.**
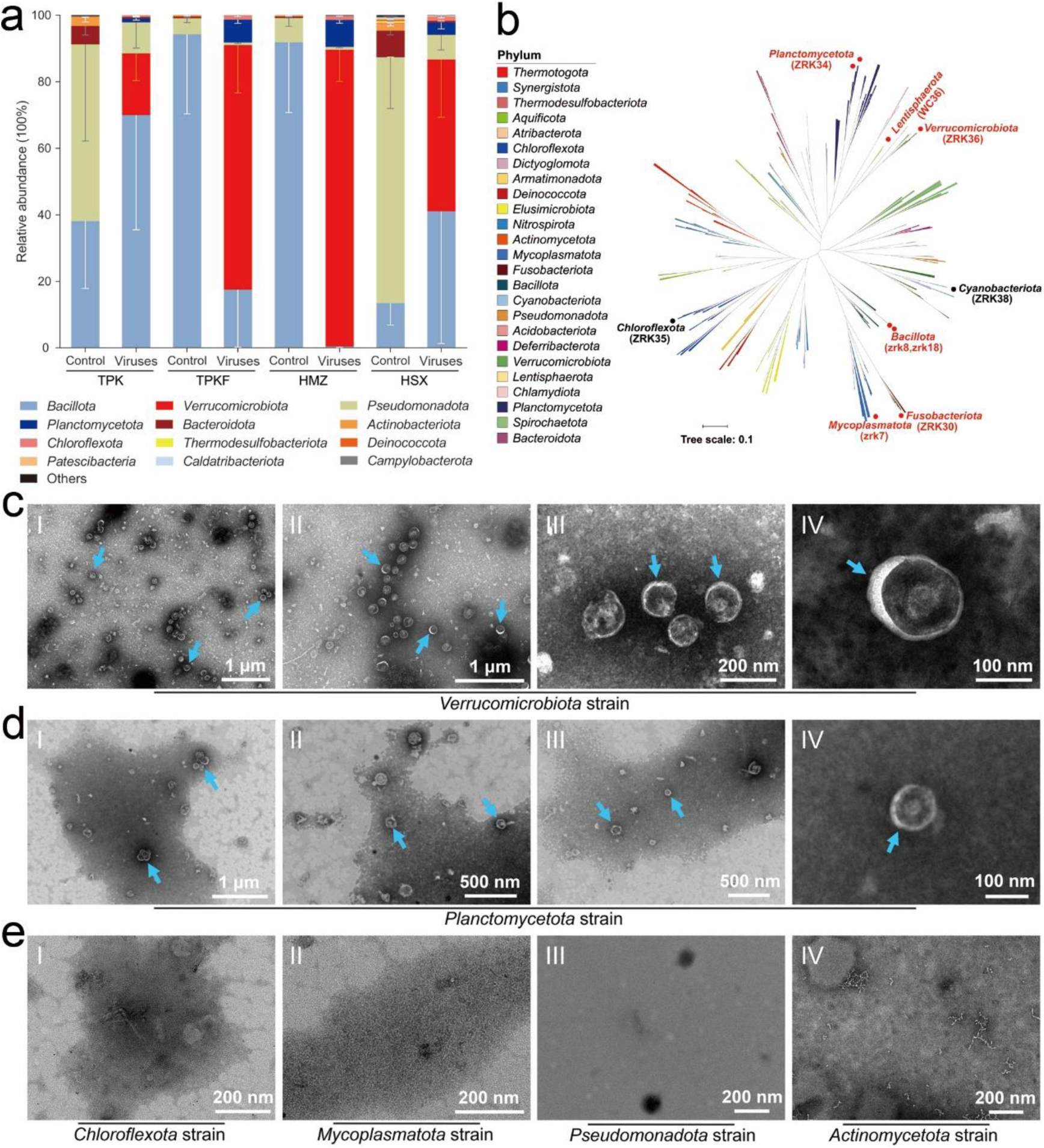
Deep-sea viruses facilitate the enrichment and cultivation of difficult-to-culture bacteria. **a**, Analysis of bacterial community structures of four deep-sea sediments (TPK, TPKF, HMZ, and HSX) in inorganic medium supplemented with or without virus suspension for 30 days. Data shown are mean ± standard deviation (n = 3 biologically independent replicates). “Control” indicates samples that were inoculated in inorganic medium supplemented with 5.0 g/L complex polysaccharides, proteins, nucleic acids, and 50 μl/mL SM buffer; “Viruses” indicates samples that were inoculated in inorganic medium supplemented with 5.0 g/L complex polysaccharides, proteins, nucleic acids, and 50 μl/mL virus suspension. **b**, Maximum-likelihood phylogenetic tree of seven novel pure cultures (red) and two enriched cultures (black) based on almost complete 16S rRNA gene sequences. Bar, 0.1 substitutions per nucleotide position. **c, d**, TEM observation of phages extracted from the supernatant of cells suspension of a *Verrucomicrobiota* strain (c) and a *Planctomycetota* strain (d) cultivated in the anaerobic 2216E medium supplemented with 3.0 g/L laminarin. Blue arrows indicated the spherical cystovirus-like phages. **e**, TEM observation of the supernatant of cells suspension of a *Chloroflexota* strain, a *Mycoplasmatota* strain, a *Pseudomonadota* strain, and an *Actinomycetota* strain that cultivated in the anaerobic 2216E medium supplemented with 3.0 g/L laminarin.

Given our previous findings that polysaccharides, specifically laminarin and starch, induce the release of chronic phages in deep-sea *Lentisphaerota* strains WC36 and zth2^26^, potentially facilitating host metabolism of these complex carbohydrates, which might be widespread among deep-sea microorganisms involved in polysaccharide degradation. To test this hypothesis, we attempted to induce phage production in our isolated deep-sea bacteria by supplementing them with polysaccharides, followed by TEM to visualize phage particles. The results showed that massive spherical cystovirus-like phages (100-150 nm in diameter, without killing the host) could be observed in supernatant concentrates of *Verrucomicrobiota* and *Planctomycetota* strains (Figs. 6c-d), but no obvious phage-like structures were observed in other bacteria (*Chloroflexota*, *Mycoplasmatota*, *Pseudomonadota*, and *Actinomycetota*) (Fig. 6e). Of note, the phyla *Lentisphaerota, Verrucomicrobiota* and *Planctomycetota* belong to the PVC superphylum^48^, and many members of this superphylum were reported to be capable of degrading and consuming polysaccharides^49–51^. Combining our results presented in this study, we speculate polysaccharides could induce the production of chronic phages in PVC bacteria and phages in turn may facilitate the metabolism of polysaccharides, which thereby promoting the host growth. It is known that members belonging to *Cyanobacteriota*, *Planctomycetota*, and *Chloroflexota* are dominant in diverse ocean environments^52^, and play crucial roles in carbon and nutrient cycling^15^. However, very few deep-sea members within these phyla have been cultured. Accordingly, they are recognized as typical difficult-to-culture bacteria. Therefore, the unique enrichment strategy associated with deep-sea viruses performed in the present study is a very novel and effective method to enrich deep-sea uncultured microorganisms, which might be developed into a common technique for isolating difficult-to-culture microbes from diverse ocean environments.

**Supplementary Fig. 2.**
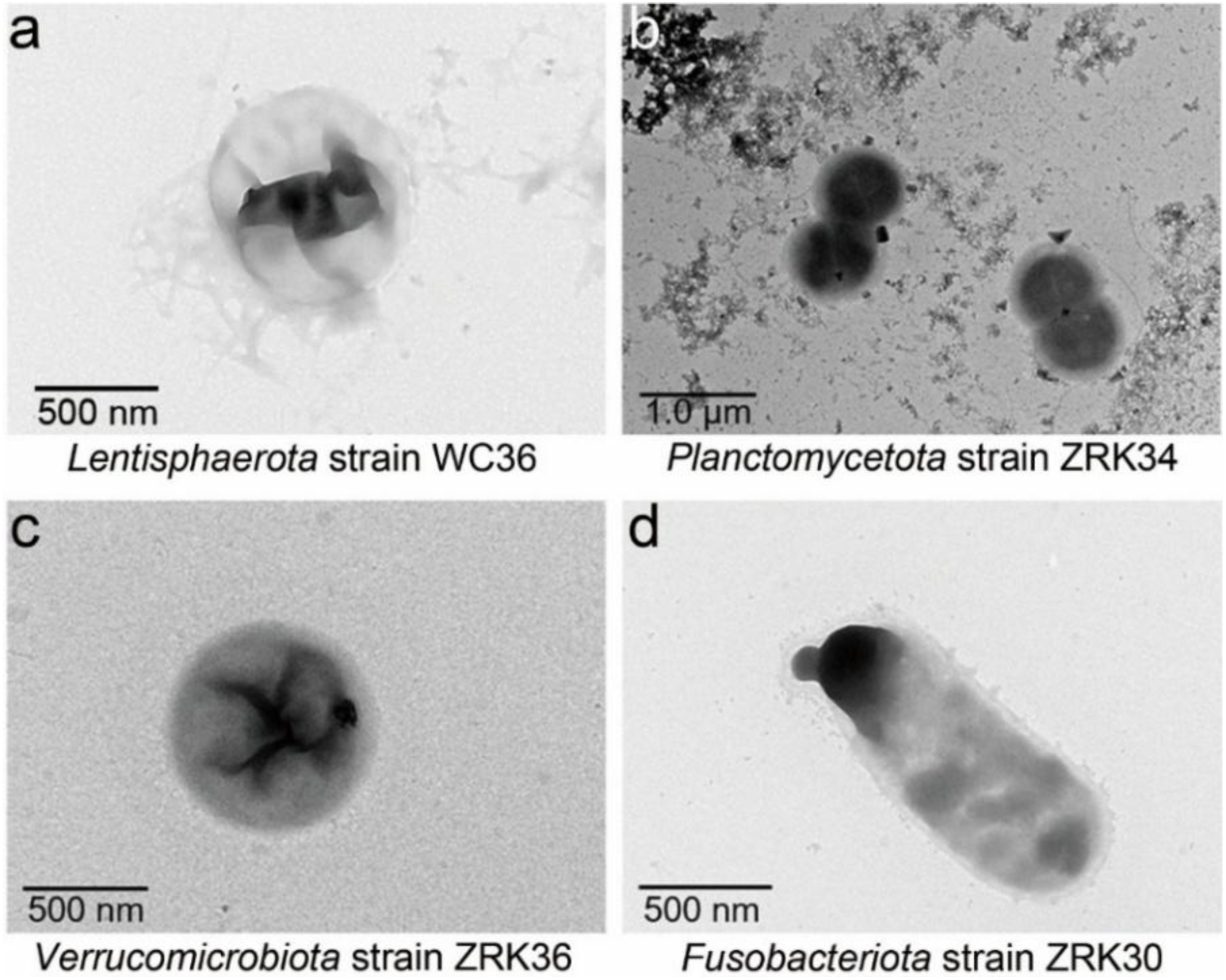
TEM observation of four deep-sea difficult-to-culture bacteria by the novel strategy involved in virus and polysaccharides. **a**, *Lentisphaerota* strain WC36. **b**, *Planctomycetota* strain ZRK34. **c**, *Verrucomicrobiota* strain ZRK36. **d**, *Fusobacteriota* strain ZRK30.

### Conclusions

In this study, we leveraged multi-omics analyses and viral enrichment experiments to comprehensively explore the diverse DNA and RNA viral communities inhabiting deep-sea sediments from distinct cold seep and seamount environments. By predicting viral host ranges, we demonstrate that both prokaryotic and eukaryotic communities are subject to viral influence, and that these interactions collectively shape the structure and function of deep-sea microbial assemblages in these critical ecosystems. Furthermore, our novel strategy of enriching for recalcitrant deep-sea microorganisms using viral fractions not only provides experimental evidence that viral-encoded functional gene can augment microbial metabolism of COM, but also enables the isolation of previously uncultured deep-sea taxa. This innovative approach offers a powerful tool for unlocking the metabolic potential of the ‘microbial dark matter’ that dominates the deep biosphere.

## Methods

### Sample collection and total organic carbon measurement

Deep-sea sediment samples from cold seep and seamount in the South China Sea were collected by *RV KEXUE* during July 2022 (Fig. 1a and Supplementary Data 1). To capture the influence of distinct deep-sea geomorphologies on microbial and viral communities, we sampled sediments from two cold-seep vents (TPK and HMZ), a site distal to cold-seep vent (TPKF), and a seamount base (HSX). The detailed geological information on collected sediments is listed in Supplementary Data 1. The total organic carbons of four deep-sea sediment samples were determined by the elemental analyzer (vario MACRO cube, Elementar, Germany) using standard methods.

### Metagenomic sequencing, assembly, and binning

Microbial DNA was extracted from these deep-sea sediment samples (TPK, TPKF, HMZ, and HSX) using the E.Z.N.A.® stool DNA Kit (Omega Bio-tek, Norcross, GA, USA) according to manufacturer’s protocols. Metagenomic shotgun sequencing libraries were constructed and sequenced at Shanghai Biozeron Biological Technology Co. Ltd. Briefly, total DNAs from these samples (20 g each) were extracted using the Qiagen DNeasy® PowerSoil® Pro Kit (Qiagen, Hilden, Germany) and the integrity of DNA was evaluated by gel electrophoresis. Then, 1.0 μg DNA was sheared by Covaris S220 Focused-ultrasonicator (Woburn, MA, USA) and sequencing libraries were prepared with a fragment length of approximately 450 bp. All samples were sequenced in the Illumina HiSeq X instrument with pair-end 150bp (PE150) mode. Raw sequence reads underwent quality trimming was performed by SOAPnuke (v1.5.6) (setting: -l 20 -q 0.2 -n 0.05 -Q 2 -d -c 0 -5 0 -7 1)^53^ and the clean data were assembled using MEGAHIT (v1.1.3) (setting:--min-count 2 --k-min 33 --k-max 83 --k-step 10)^54^. Assemblies of these samples were automatically binned using Maxbin2 (version 2.2.7, -markerset 40 option)^55^, metaBAT2 (version 2.12.1, -m 1500 and --unbinned parameters)^56^ and Concoct (version 0.4.0, default settings)^57^. MetaWRAP (version 1.3.2, default parameters)^58^ was used to purify and organize data to generate the final bins. Finally, the completeness and contamination of metagenome-assembled genomes (MAGs) were assessed by the checkM (v1.0.18)^59^.

### Phylogeny and abundance analysis of deep-sea MAGs

Genome taxonomy of MAGs from the four deep-sea sediment samples was determined using GTDB-Tk v2.1.0^60^. A maximum-likelihood phylogeny of the 65 MAGs, along with 13 downloaded NCBI referenced genome sequences, was inferred using IQ-TREE v2.2.0^61^ with the GTR+F+I+G4 model based on a concatenation of 120 bacterial or 122 archaeal marker genes identified by GTDB-Tk v2.1.0; the resulting tree was visualized using the online tool Interactive Tree of Life (iTOL v5)^62^. MAG relative abundances were determined by mapping clean reads to MAGs using CoverM v0.6.1 (parameters: -contig -mrpkm –trim-min 5 –trim-max 95) to calculate RPKM values (Supplementary Data 2).

### Virus purification

To explore the abundance and genomic composition of DNA and RNA viruses in deep-sea environments, four deep-sea sediment samples (TPK, TPKF, HMZ, and HSX) were used to enrich and isolate viruses. Briefly, 20 g deep-sea sediment sample mixed with 8 mL SM buffer and 20 sterilized glass beads were put into a 50 mL centrifuge tube, then vibrated and mixed at 4 °C for 30 min. Next, the mixture was centrifuged at 5,000 ×*g* for 10 min at 4 °C, and the supernatant was transferred to a new 50 mL centrifuge tube. The preceding steps were repeated for four times. After that, all supernatants were collected and centrifuged at 10,000 × *g* for 30 min at 4 °C to remove impurities. Meanwhile, supernatants containing viruses were filtered with a 0.22 μm filter membrane and then precipitated with 10% PEG8000 at 4 °C for 12 h. Finally, the precipitated supernatants were centrifuged at 200,000 ×*g* for 2 h at 4 °C, and the pellet containing phage-particles was obtained. After resuspended with the SM buffer (0.01% gelatin, 50 mM Tris-HCl, 100 mM NaCl and 10 mM MgSO_4_), the virus suspension was used for viral metagenomic sequencing and Transmission electron microscopy (TEM) observation.

### Transmission electron microscopic observation

To observe the morphology of bacteriophages, phage virions were allowed to adsorb to the copper grid for 20 min, and then stained with the phosphotungstic acid for 30 s. Next, micrographs were taken with TEM (HT7700, Hitachi, Japan) with a JEOL JEM 12000 EX (equipped with a field emission gun) at 100 kV. To observe the morphology of pure deep-sea bacterial cultures, its cell suspension incubated in the anaerobic 2216E medium supplemented with 3.0 g/L laminarin was centrifuged at 5,000 ×*g* for 10 min to obtain cell pellets. Subsequently, one part of the cell collection was adsorbed to the copper grids for 20 min, then washed with 10 mM phosphate buffer solution (PBS, pH 7.4) for 10 min and dried at room temperature for TEM observation.

### Viral metagenomic sequencing

For DNA viral metagenomic sequencing, total DNAs were extracted from the purified virus samples using the E.Z.N.A.® stool DNA Kit (Omega Bio-tek, USA) according to manufacturer’s protocols. High-quality DNA sample (OD260/280 = 1.8∼2.2, OD260/230 ≥2.0) was used to construct sequencing library. DNA viral metagenomic libraries were prepared following TruSeqTM Nano DNA sample preparation Kit from Illumina (San Diego, CA), using 1.0 ug of total DNA. DNA end repair, A-base addition and ligation of the NGS-indexed adaptors were performed according to NGS’s protocol. Libraries were then size selected for DNA target fragments of ∼400bp on 2% Low Range Ultra Agarose followed by PCR amplified using Phusion DNA polymerase (NEB, USA) for 15 PCR cycles.

For RNA viral metagenomic sequencing, total RNA was extracted from the purified virus samples using TRIzol® Reagent according to the manufacturer’s instructions (Invitrogen) and genomic DNA was removed using DNase I (TaKara, Japan). Then RNA quality was determined using 2100 Bioanalyser (Agilent) and quantified using the ND-2000 (NanoDrop Technologies). High-quality RNA sample (OD260/280 = 1.8∼2.2, OD260/230 ≥2.0) was used to construct sequencing library. RNA viral metagenomic libraries were prepared following TruSeqTM RNA sample preparation Kit (Illumina, USA), using 5.0 μg of total RNA. Shortly, rRNA removal by Ribo-ZeroTM rRNA Removal Kits (Epicenter, Sweden), fragmented using fragmentation buffer. cDNA synthesis, end repair, A-base addition and ligation of the NGS-indexed adaptors were performed according to NGS’s protocol. Libraries were then size selected for cDNA target fragments of 200-300 bp on 2% Low Range Ultra Agarose followed by PCR amplified using Phusion DNA polymerase (NEB) for 15 PCR cycles. DNA and RNA metagenomic sequencing was performed by Shanghai Biozeron Biotechnology Co., Ltd. (Shanghai, China). All samples were sequenced in the next generation sequencing platform with pair-end 150bp (PE150) mode. The raw paired end reads were trimmed and quality controlled by Trimmomatic with parameters (SLIDINGWINDOW:4:15 MINLEN:75) (version 0.40, http://www.usadellab.org/cms/?page=trimmomatic)^63^. Then clean reads that aligned to the prokaryotic genomes were also removed. This set of high-quality reads was then used for further analysis.

### Viral metagenomic *de novo* assembly, gene prediction and annotation

Clean sequence reads were generated a set of contigs of each sample using MegaHit with “--min-contig-len 500” parameters^54^. For the assembled genome, the above 5000 bp (DNA) and 2000 bp (RNA) assembly sequences were selected, and those sequences were performed virus identification using VirFinder (parameters: score > 0.7, *P* value < 0.05)^64^, VirSorter2^65^ and IMG/VR (Integrated Microbial Genomes, https://img.jgi.doe.gov/cgi-bin/vr/main.cgi) database. vOTUs (viral operational taxonomic units) were clustered using Mummer software to compare candidate virus sequences, with greater than 95% similarity and 85% coverage of the total sequence length and the longest representative one within each cluster was considered as vOTUs^66^. All the genes within vOTUs were predicted by METAProdigal (http://compbio.ornl.gov/prodigal/)^67^ and annotated by BLASTX against the NCBI non-redundant (NR), String, eggNOG, and KEGG databases (E-value < 1.0×10^−5^, Similarity >= 30%). Gene Ontology (GO) annotations for unique assembled transcripts were obtained using BLAST2GO (http://www.blast2go.com/b2ghome) to characterize biological processes, molecular functions, and cellular components. Metabolic pathway analysis was performed using the Kyoto Encyclopedia of Genes and Genomes (KEGG, http://www.genome.jp/kegg/).

### Viral taxonomic assignments and abundance profiles

Open reading frames (ORFs) were predicted from vOTU sequences using Prodigal v2.6.3^68^ (-p meta -g 11 -m -c) and classifying vOTUs against the NCBI viral RefSeq database (2023-04-26) using CAT v5.0.3^69^ with the Last Common Ancestor algorithm. vOTU relative abundances were determined by mapping clean reads to vOTUs using CoverM v0.6.1 (contig and genome modes, -m rpkm --trim-min 5 - -trim-max 95) to calculate RPKM (reads per kilobase per million mapped reads) values.

### Virus–host prediction

Virus-host linkages were predicted using four in silico strategies: (1) Nucleotide sequence homology. Sequences of vOTUs and prokaryotic MAGs were compared using BLASTn (≥75% coverage, ≥70% nucleotide identity, ≥50 bit score, E-value ≤ 0.001). (2) Oligonucleotide frequency (ONF). VirHostMatcher v1.0 (with default parameters)^70^ was used to identify linkages based on oligonucleotide frequency (d2*≤0.2, as a match). (3) Transfer RNA (tRNA) match. The tRNAs from prokaryotic MAGs and vOTUs were identified using ARAGORN v1.2.65^71^ (-t option), requiring ≥90% length identity across ≥90% of sequences by BLASTn^72^. (4) CRISPR spacer match. CRISPR arrays were assembled from quality-controlled reads using Crass v1.0.1 with default parameters^73^. Spacers were matched against viral contigs (≤1 mismatch) and repeats from the same CRISPR array were compared against prokaryotic MAGs using BLASTn, creating virus-host links. Putative microbial hosts with adjacent cas genes, identified using MetaErg v1.2.2^74^, were considered high-confidence linkages. When multiple hosts were predicted for a vOTU, the linkage supported by multiple approaches was prioritized. Otherwise, to assign a single host, linkages were ranked as follows: (1) CRISPR spacer match with adjacent cas genes; (2) CRISPR spacer match without adjacent cas genes; (3) tRNA match or nucleotide sequence homology; (4) ONF comparison. The highest-ranked host was assigned^20^.

### Viral life strategy prediction

Viral life strategies were predicted using VIBRANT v1.2.0^75^ and CheckV v0.9.0^76^. Temperate strategies were inferred by identifying vOTUs containing provirus integration sites or integrase genes. vOTUs identified through this pipeline were classified as temperate; remaining vOTUs were designated as unknown.

### Metatranscriptomic analysis

To explore the actual metabolic characteristics of deep-sea DNA and RNA viruses conducted in the deep-sea environments, the metatranscriptomic analysis was performed. These same sediment samples (TPK, TPKF, HMZ, and HSX) were used for metatranscriptomic sequencing analysis in Shanghai Biozeron Biothchnology Co., Ltd. (Shanghai, China). Total RNAs were extracted from these sediments using TRIzol® Reagent according the manufacturer’s instructions and genomic DNA was removed using DNase I (TaKara, Japan). Then RNA quality was determined using 2100 Bioanalyser (Agilent) and quantified using the ND-2000 (NanoDrop Technologies). High-quality RNA sample (OD260/280 = 1.8∼2.2, OD260/230 ≥2.0, RIN ≥6.5, 28S:18S ≥1.0, >10 μg) is used to construct sequencing library. Metatranscriptomic libraries were prepared following TruSeq TM Stranded Total RNA Sample Preparation Kit from Illumina (San Diego, CA, USA), using 5 μg of total RNA. Briefly, rRNA removal by Ribo-Zero TM rRNA Removal Kits from Illumina (San Diego, CA, USA), fragmented using fragmentation buffer. cDNA synthesis, end repair, A-base addition, and ligation of the Illumina-indexed adaptors were performed according to Illumina’s protocol. Libraries were then size selected for cDNA target fragments of 200-300 bp on 2% Low Range Ultra Agarose followed by PCR amplified using Phusion DNA polymerase (NEB, USA) for 15 PCR cycles. All samples were sequenced in the Illumina HiSeq 2500 instrument. Libraries were prepared with a fragment length of approximately 450 bp. Paired-end reads were generated with 150 bp in the forward and reverse directions. Raw metatranscriptomic reads were quality filtered in the same manner as metagenomes. Subsequently, these high-quality metatranscriptomic reads were mapped to the predicted protein-coding genes from all the DNA and RNA viruses using Salmon v.1.5.0^77^ in mapping based mode (parameters: -validateMappings -meta). Transcript levels of functional genes involved in the degradation of COM within DNA and RNA viruses were normalized to transcripts per million (TPM).

### Operational taxonomic units (OTUs) analysis

Since deep-sea DNA and RNA viruses may have the ability to assist bacterial hosts in metabolizing macromolecules, we designed unique enrichment media (inorganic medium supplemented with 5.0 g/L complex polysaccharides, proteins, nucleic acids, and 50 μl/mL SM buffer, as the control group; inorganic medium supplemented with 5.0 g/L complex polysaccharides, proteins, nucleic acids and 50 μl/mL virus suspension, as the experimental group) to enrich and culture deep-sea microorganisms. Specifically, the four selected sediment samples (TPK, TPKF, HMZ, and HSX) were cultured at 28 °C for one month in the enrichment media as described above. Each medium had three biologically independent replicates. To understand the bacterial community structure of these enrichment cultures, 2 mL of each enrichment medium was harvested and sent to Novogene (Tianjin, China) for OTUs sequencing. Briefly, total DNAs from different samples were extracted respectively by the CTAB/SDS method^78^ and diluted to 1 ng/µL with sterile water, which were used for PCR templates. 16S rRNA genes of distinct regions (16S V3/V4) were amplified using specific primers (341F: 5’-CCTAYGGGRBGCASCAG-3’ and 806R: 5’-GGACTACNNGGGTATCTAAT-3’). The PCR products were purified with a Qiagen Gel Extraction Kit (Qiagen, Germany) following the manufacturer’s instructions for libraries construction. Sequencing libraries were generated using TruSeq® DNA PCR-Free Sample Preparation Kit (Illumina, USA) according to the manufacturer’s instructions. The library quality was assessed on the Qubit@ 2.0 Fluorometer (Thermo Scientific, USA) and Agilent Bioanalyzer 2100 system. Then, the library was sequenced on an Illumina NovaSeq platform and 250 bp paired-end reads were generated. Paired-end reads were merged using FLASH (V1.2.7, http://ccb.jhu.edu/software/FLASH/)^79^, which was designed to merge paired-end reads when at least some of the reads overlap with those generated from the opposite end of the same DNA fragments, and the splicing sequences were called raw tags. Quality filtering on the raw tags was performed under specific filtering conditions to obtain the high-quality clean tags^80^ according to the QIIME (V1.9.1, http://qiime.org/scripts/split_libraries_fastq.html) following standard protocols. The tags were compared with the reference database (Silva database, https://www.arb-silva.de/) using UCHIME algorithm (UCHIME Algorithm, http://www.drive5.com/usearch/manual/uchime_algo.html)^81^ to detect chimera sequences, and then the chimera sequences were removed^82^. Lastly, Uparse software (Uparse v7.0.1001, http://drive5.com/uparse/)^83^ was used to analyze these sequences. Sequences with ≥97% similarity were assigned to the same OTUs. The representative sequence for each OTU was screened for further annotation. For each representative sequence, the Silva Database (http://www.arb-silva.de/)^84^ was used to annotate taxonomic information based on Mothur algorithm.

### Phylogenetic analysis of pure-cultured deep-sea bacteria

The maximum likelihood phylogenetic tree of the novel pure cultures (strains WC36, ZRK34, ZRK36, ZRK30, zrk7, zrk8, and zrk18) and two enriched cultures (respectively predominant by strains ZRK35 and ZRK38) was based on almost complete 16S rRNA gene sequences. The 16S rRNA gene sequences of other phyla bacteria were all obtained from the NCBI databases. All the sequences were aligned by MAFFT version 7^85^ and manually corrected. The phylogenetic trees were constructed using the W-IQ-TREE web server (http://iqtree.cibiv.univie.ac.at)^86^. Finally, we used the online tool Interactive Tree of Life (iTOL v5)^62^ to edit the tree.

### Observation of bacteriophages induced by the supplement of polysaccharide

To investigate whether polysaccharides could induce phage production in deep-sea bacteria, we selected cultured representatives from six phyla (*Verrucomicrobiota*, *Planctomycetota*, *Chloroflexota*, *Mycoplasmatota*, *Pseudomonadota*, and *Actinomycetota*) in our laboratory. All strains were cultivated in the anaerobic liquid 2216E medium supplemented without or with 3 g/L laminarin at 28 °C for 10 days. Subsequently, we performed the phage isolation and TEM observation as described above.

## Data availability

The raw sequencing reads from the DNA and RNA viral metagenomes have been deposited to NCBI Short Read Archive (accession number: PRJNA1230403 and PRJNA1230580). The raw metatranscriptomic sequencing data have been deposited to NCBI Short Read Archive (accession number: PRJNA1230164). The raw amplicon sequencing data have also been deposited to NCBI Short Read Archive (accession number: PRJNA1230549). The whole 16S rRNA gene sequences of strains WC36, ZRK34, ZRK36, ZRK30, zrk7, zrk8, zrk18, ZRK35, and ZRK38 have been deposited in the GenBank database with accession numbers OK614042, MT076070, MZ646149, OM892919, OP108612, MW581533, MW647584, MW581537, and OP108622, respectively.

## ACKNOWLEDGEMENTS

This work was supported by the National Natural Science Foundation of China (Grant No. 42406104), the NSFC Innovative Group Grant (No. 42221005), Science and Technology Innovation Project of Laoshan Laboratory (Grant Nos. LSKJ202203103 and 2022QNLM030004-3), Shandong Provincial Natural Science Foundation (Grant Nos. ZR2024ZD49, ZR2021ZD28 and ZR2023QD010), Major Research Plan of the National Natural Science Foundation (Grant No. 92351301), and Taishan Scholars Program (Grant Nos. tstp20230637 and tsqn202312264). We extend special appreciation to the captain and crew of the *RV KEXUE*, as well as the FAXIAN ROV team for assistance with sample collection.

## AUTHOR CONTRIBUTIONS

CW and CS conceived and designed the study; CW conducted most of the experiments; CW and RZ collaboratively analyzed the omics data; CW, RZ, and CS lead the writing of the manuscript; all authors contributed to and reviewed the manuscript.

